# Deep ubiquitination site profiling by single-shot data-independent acquisition mass spectrometry

**DOI:** 10.1101/2020.07.23.218651

**Authors:** Martin Steger, Phillip Ihmor, Mattias Backman, Stefan Müller, Henrik Daub

## Abstract

We report a highly optimized proteomics method for in-depth ubiquitination profiling, which combines efficient protein extraction and data-independent acquisition mass spectrometry (DIA-MS). Employing DIA for both spectral library generation and single-shot sample analysis, we quantify up to 70,000 ubiquitinated peptides per MS run with high precision, data completeness and throughput. Our approach resolves the dynamics of ubiquitination and protein degradation with an unprecedented analytical depth.

---

The ubiquitin proteasome system (UPS) regulates a plethora of intracellular processes through the interplay of ubiquitin ligases and deubiquitinating enzymes (DUBs). Ubiquitin is predominantly attached to lysine (K) residues on substrate proteins and itself can be further ubiquitinated at multiple sites to form branched polymeric chain structures, which encode for specific signals such as proteasomal degradation^1^.

MS-based proteomics has emerged as powerful tool to investigate global cellular ubiquitination, with enormous potential to profile UPS-targeting drugs^2^. Current ubiquitinomics workflows rely on antibodies to purify K-GG or K-GGRLRLVLHLTSE remnant peptides, which result from digesting ubiquitin-modified proteins with trypsin or Lys-C, respectively^3,4^. While tens of thousands of sites have been mapped by label-free MS, large-scale ubiquitination profiling requires high protein input and extensive peptide fractionation, limiting its application to small sample numbers. Isobaric peptide labeling strategies can partially address these limitations^5^; however, they typically suffer from lower ubiquitination site coverage and less accurate quantification, further confounded by labeling batch effects.

To overcome these limitations, we first re-evaluated the urea-based lysis procedure for ubiquitinomics^6^. Since sodium deoxycholate (SDC)-based protein extraction is best suited for proteome analyses^7,8^, we tested SDC buffer instead of urea. We tryptically digested 2 mg of total protein from MG-132-treated HCT116 cells and enriched K-GG remnant peptides, followed by 125 min single-shot LC-MS analysis in data-dependent acquisition (DDA) mode. We reasoned that immediate boiling of the lysates and the high concentration of DUB-inactivating chloroacetamide in the SDC buffer would increase ubiquitin site coverage. Indeed, compared to urea, SDC lysis yielded on average 27% more K-GG peptides per sample (∼29,000 vs. 21,000) while preserving relative enrichment specificity (Fig. 1, Supplementary Fig. 1 and Supplementary Table 1).

**Fig. 1.**
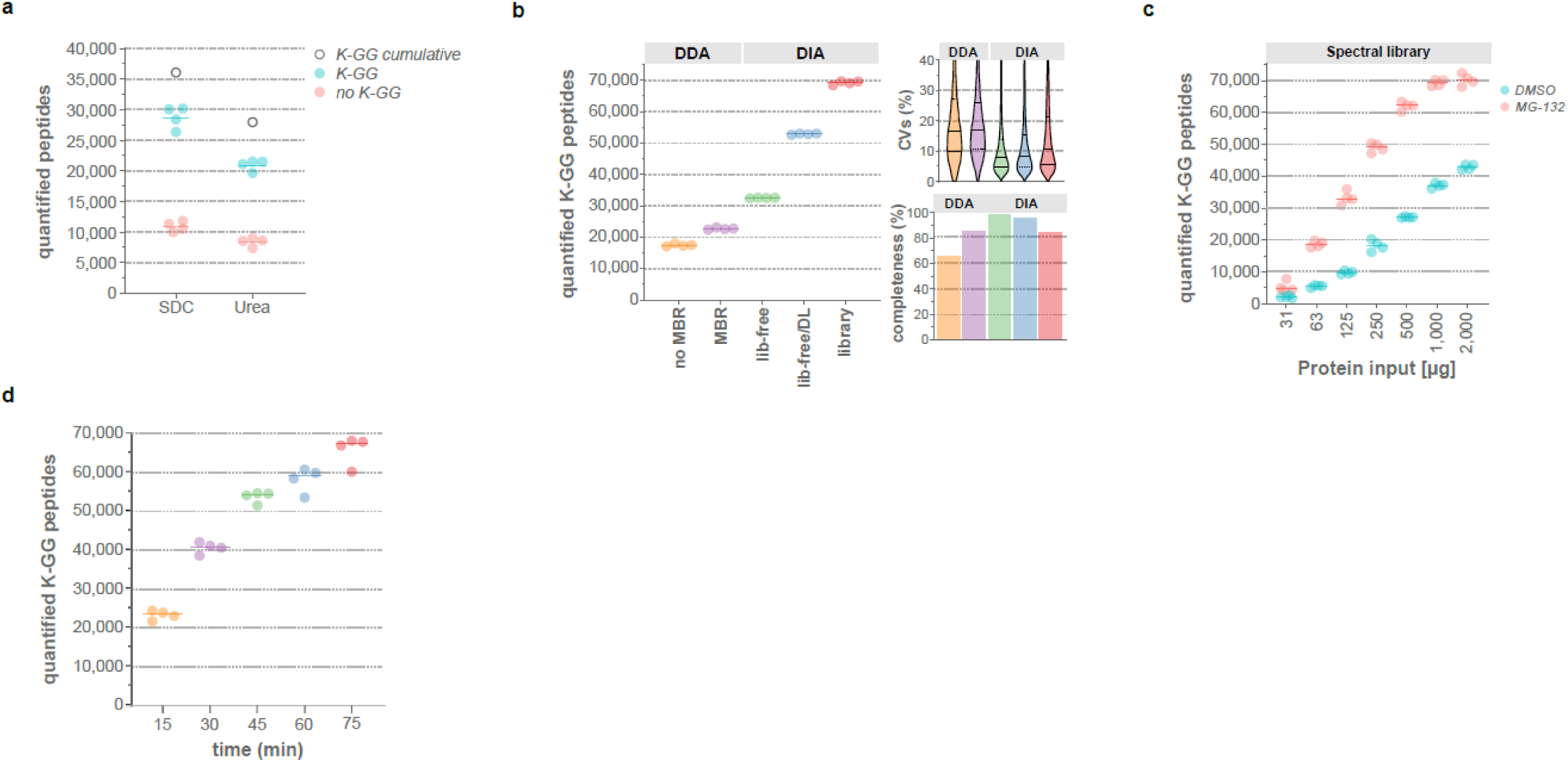
Implementation and optimization of deep coverage, single-shot MS-based ubiquitinomics workflow. **a**, Quantified K-GG peptides and non -K-GG peptides detected in 125 min DDA LC-MS runs upon cell lysis with either urea or SDC lysis buffer. 2 mg of total protein of HCT116 cells were used for each replicate (only half of the sample was injected). Open circles show cumulative numbers of K-GG peptide identifications from four replicates. **b**, Quantified K-GG peptides with 2 mg of input per replicate in a side-by-side comparison of different DDA and DIA MS and data analysis methods (MBR= match between runs, lib-free = library-free, DL= deep learning) based on replicate 75 min single-shot LC-MS runs. Library-based DIA data analysis was performed with an ultra-deep K-GG peptide library generated by library-free DIA analysis, as described in detail in *Methods*. Coefficients of variation (CVs) and data completeness (percentage of valid values in the data matrix) for each method are shown. For the CVs, continuous lines demarcate the median and dashed lines the upper and lower quartiles. **c**, HCT116 cells were cultured in the presence or absence of MG-132 (10 µM, 6 h) and K-GG peptides from different protein inputs quantified by single-run DIA-MS using the ultra-deep K-GG spectral library. **d**, Number of K-GG peptides quantified with DIA using different LC gradient lengths. The same spectral library as in **b and c** was used for data processing.

Even though we quantified more than 25,000 K-GG peptides by single-shot MS, due to the semi-stochastic sampling inherent to DDA, only 40-50% of those identifications were without missing values in replicate samples. Therefore, we evaluated data-independent acquisition (DIA), as it was reported to outperform DDA in terms of quantification precision, number of identifications, and data completeness^9^. Using the library-free search option of the DIA-NN^10^ software, we initially compared 75 and 125 min single-shot DIA methods, identifying just 18% less ubiquitinated peptides with the 75 min method, while reducing acquisition time by 40% (Supplementary Fig. 2). We therefore employed 75 min gradients for subsequent direct comparison of DIA to DDA. We measured four injection replicates of K-GG enriched peptides with both acquisition modes and analyzed the resulting DDA and DIA raw files with MaxQuant and DIA-NN, respectively^10,11^. For DIA-NN, we first tested library-free analysis, with or without deep learning (Supplementary Fig. 3). Strikingly, compared to DDA, library-free searching without deep learning resulted in 30% more K-GG peptides on average, with overall less than 2% of missing values and a median CV< 8%. IDs further increased by an additional 38% when enabling deep learning, resulting in as many as 53,000 quantified K-GG peptides per replicate (Fig. 1b and Supplementary Fig. 4). For spectral library generation, we prepared 12 high pH reversed-phase fractions of a tryptic HCT116 digest and enriched each fraction for K-GG peptides. Encouraged by our single-shot data, we compared both conventional DDA and library-free DIA analyses for K-GG peptide spectral library generation (see Methods). Notably, the resulting DIA library contained 15% more K-GG peptides than its DDA counterpart (108,427 vs. 92,317) (Supplementary Fig. 3). By analyzing another set of fractionated K-GG peptide samples, we increased DIA library size to 131,559 K-GG peptides. Harboring 111,707 distinct ubiquitination sites mapping onto 11,983 proteins, this dataset contains more entries than all previous ubiquitinomics studies combined (110,072 in PhosphoSitePlus v6.5.9.2) (Supplementary Table 2). Using this library for single-shot DIA analysis, we quantified on an average 69,231 K-GG peptides in 75 min MS runs, with high data completeness and a median CV of about 10% (Fig. 1b, Supplementary Fig. 4 and Supplementary Table 3).

Previous ubiquitinomics studies used proteasome inhibitors to block protein degradation, thus preserving and boosting protein ubiquitination^3,4^. However, such compounds are cytotoxic and result in an accumulation of newly synthesized, misfolded proteins, thus perturbing global protein turnover^12^. Furthermore, proteasomal inhibition hampers the monitoring of protein degradation as a direct consequence of ubiquitination. To test whether our library-based method could overcome these limitations even with low sample amounts, we applied it to different input amounts of both MG-132- and DMSO-treated cell lysates. While we reached saturation of about 70,000 K-GG peptide identifications with 1 mg protein input from MG-132-treated cells, we observed a steady increase to about 40,000 with 2 mg input in DMSO-treated samples (Fig. 1c and Supplementary Table 4). Notably, with just 500 µg of protein, we still quantified more than 30,000 K-GG peptides in untreated cells, highlighting the feasibility of in-depth ubiquitinomics for unperturbed samples available in limited amounts.

To evaluate the potential for higher throughput applications, we tested shorter LC gradients in combination with our optimized K-GG enrichment protocol. Even a 15 min gradient consistently quantified more than 20,000 K-GG peptides with good precision, highlighting that our method can achieve significantly higher coverage than TMT-based multiplexing protocols of similar throughput^5^ (Fig. 1d, Supplementary Fig. 5 and Supplementary Table 4).

Currently, there are six clinically approved drugs targeting the UPS and many more are in preclinical testing^2^. However, the modulation and time-resolved interplay of protein ubiquitination and degradation upon treatment with such drugs has been challenging to analyze on a truly proteome-wide scale. To evaluate our DIA-MS workflow for this application, we performed a time-course experiment with the selective Usp7 inhibitor FT671^13^ (Supplementary Fig. 6). Usp7 is of particular interest because of its role in deubiquitinating and stabilizing the E3 ligase Mdm2. As Mdm2 promotes p53 degradation, inhibition of Usp7 leads to p53-dependent tumor growth suppression^14,15^. However, as recently developed Usp7 inhibitors only partially block tumor growth and other Usp7 targets have been reported, the underlying ubiquitin signaling network may be more complex than initially thought^13,15–17^. We tested five different FT671 treatment times (between 15 min and 6 h) without proteasome inhibition, and measured both ubiquitinome and proteome in single-shot mode. Employing HCT116-derived spectral libraries, we quantified about 40,000 K-GG peptides and 10,000 proteins in each sample (Supplementary Fig. 6). Although DIA generally features high data completeness, low abundant Usp7 target sites below the detection limit in untreated cells are potentially missed and excluded from statistical analysis, even if induced and robustly quantified upon Usp7 inhibition. Therefore, we devised and validated a missing value rescue strategy, thus obtaining complete, quantitative data matrices for 42,866 K-GG peptides and 10,302 proteins, including 7,374 proteins for which we mapped at least one ubiquitination site (Supplementary Fig. 7 and Supplementary Table 6).

Already at 15 min of FT671 treatment, we found 1,277 ubiquitinated peptides significantly up-regulated by more than two-fold, which mapped to 599 proteins (Supplementary Fig. 8). Strikingly, only 37 of those rapidly ubiquitinated proteins were significantly downregulated by more than 20% over the 6 h time course. Interestingly, enrichment analysis revealed that these proteins are significantly overrepresented in transcriptional regulation and the ubiquitin conjugation pathway (Fig. 2a, Supplementary Fig. 9).

**Fig. 2.**
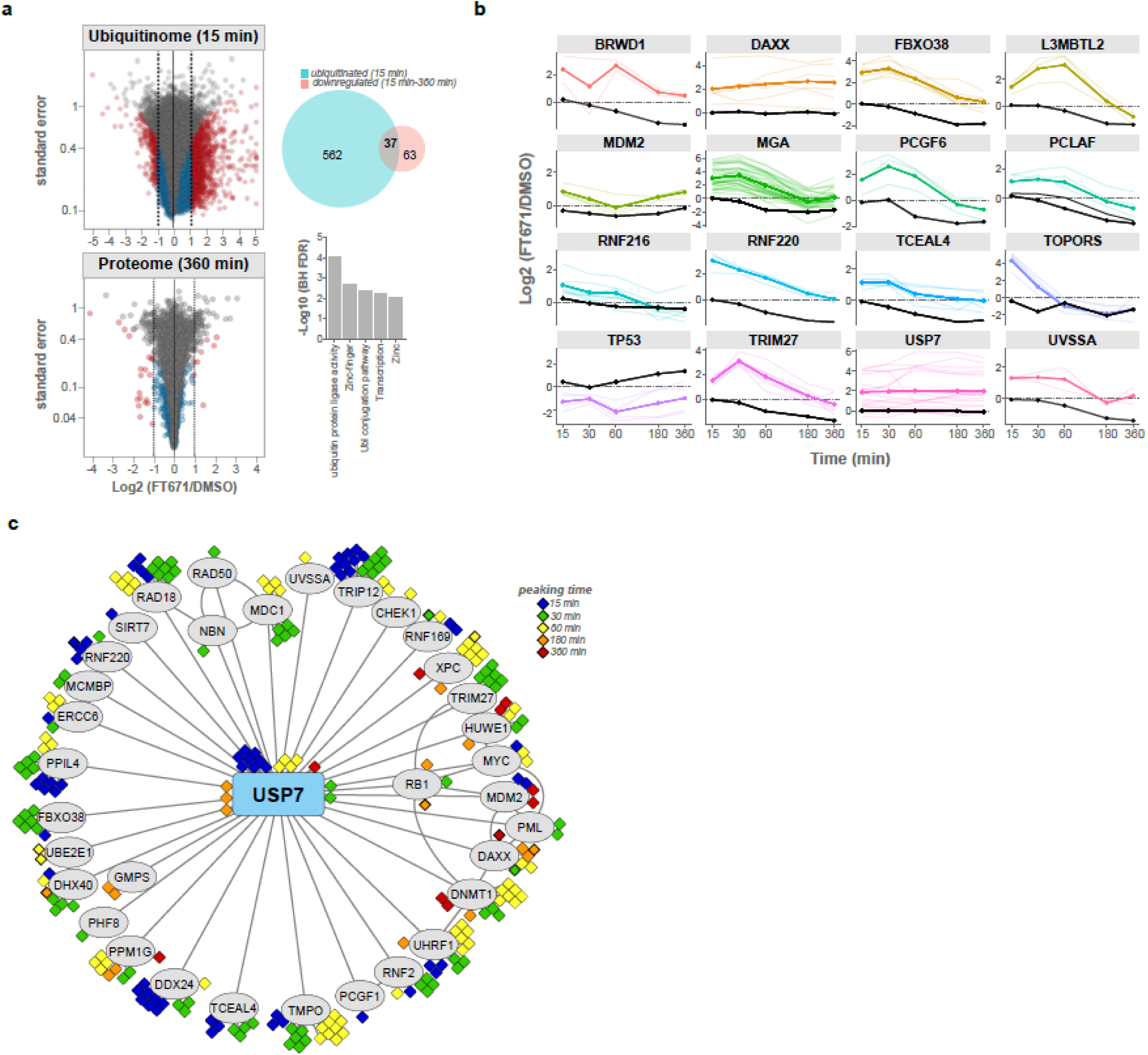
Quantification of both the ubiquitinome and proteome in a FT671 time course study. **a**, Volcano plots of the ubiquitinome (15 min) and the proteome (6 h) after FT671 treatment are shown on the left. The Venn diagram shows significantly and >2-fold ubiquitinated proteins at 15 min (turquoise) and proteins with detected ubiquitination sites that were significantly downregulated by more than 20% over the 6 h time course (pink). The overlapping proteins were significantly enriched for transcriptional regulators and E3 ubiquitin ligases **b**, Profile plots of selected targets showing MS-based quantifications of the proteins (in black) and their corresponding ubiquitination sites (colored), which were significantly and >2-fold regulated at 15 min of FT671 treatment. Both average (solid lines) and individual (transparent lines) ubiquitinated peptide profiles are plotted. **c**, Significantly and more than two-fold upregulated ubiquitinated peptides at 15 min of FT671 treatment were mapped onto a BioGrid^20^ network for reported Usp7 interacting proteins. The peak times of the individual ubiquitination sites are color-coded.

We detected upregulated ubiquitination sites for many reported Usp7 targets, including Trim27, UvssA and Mdm2^14,16,17^. For some of those, the initial increase of one or more ubiquitination sites led to the degradation of the protein with varying kinetics at later time points (Fig. 2b). Although four Mdm2 sites were induced already after 15 min, its protein levels only transiently decreased to return to baseline thereafter, likely due to transcriptional upregulation mediated by p53. Concomitantly, ubiquitination sites on p53 initially decreased in intensity, resulting in protein stabilization at later time points (Fig. 2b). Besides known substrates, we found a number of degraded proteins without any reported connection to Usp7, including ubiquitin ligases (Rnf220, Rnf216, Topors) and various transcriptional regulators (L3mbtl2, Pcgf6, Tceal4). Interestingly, many proteins, including the Usp7 interaction partner Daxx^18^, were not degraded, even though multiple sites were strongly upregulated throughout the time course (Fig. 2b). This highlights that our time-resolved quantification of both the proteome and the ubiquitinome in the absence of proteasome inhibition distinguishes ubquitination triggering apparent protein degradation from other types of ubiquitin signals.

To pinpoint primary Usp7 targets and to get more insights into Usp7-mediated signaling, we mapped our ubiquitinomics data onto a Usp7 interaction network retrieved from BioGrid^19^. Strikingly, we identified significantly upregulated sites at 15 min of FT671 treatment on 62% of all reported physical Usp7 interactors, strengthening the evidence that these proteins represent direct Usp7 targets (Fig. 2c).

In conclusion, our optimized ubiquitinomics workflow precisely and consistently quantifies tens of thousands of ubiquitinated peptides at high throughput. We envision that integration of deep ubiquitinome and proteome data will allow dissecting ubiquitin signaling of key biological processes.

## Acknowledgements

We thank all members of Evotec München GmbH and especially S. Blencke, A. Kirgis, T. Lanzinner, J. Linnemann, S. Marx, J. Modrow and D. Turnblad-Phillips for technical assistance. M. Bader, C. Kluger, B. Kracher and M. Segura-Lepe for help in developing the DIA data analysis pipeline. U. Ohmayer for help in estimating the FDR and V. Demichev for helpful discussions around the DIA-NN software.

## Author contributions

M.S. conceived the study. H.D., S.M. and M.S. designed the experiments. M.B. and P.I. and M.S. analyzed the data. All authors interpreted the data. M.S. coordinated studies and H.D. and M.S. wrote the manuscript. All authors read and approved the final manuscript.

## Author information

Correspondence and requests for materials should be addressed to

## Materials and Methods Reagents

MG-132 (474787), 2-chloroacetamide (CAA, 22790), tris(2-carboxyethyl)phosphine hydrochloride (TCEP, C4706), sodium deoxycholate (SDC, 30970), dimethyl pimelimidate dihydrochloride (DMP, D8399), ethanolamine (411000), Na_2_HPO_4_ (S9763), 3-(N-morpholino)propanesulfonic acid (MOPS, M5162), sodium azide (S2002) and sodium tetraborate decahydrate (S9640) were from Millipore Sigma. Ethylenediaminetetraacetic acid (EDTA, 108418), Tris(hydroxymethyl)-aminomethan (Tris, 108382), trifluoracetic acid (TFA, 108178), ammonium hydroxide (NH_4_OH, 533003), sodium chloride (106404), ethyl acetate (103649), acetonitrile (ACN, 100030) from Merck. Formic acid (FA, 56302) from Fluka. FT671 from AOBIUS. Trypsin/LysC mix (V5071 or V5072) from Promega. PTMScan® Ubiquitin Remnant Motif (K-ε-GG) Kit (#5562) from Cell Signaling Technology.

### Cell culture, drug treatments and cell lysis

HCT116 were purchased from ATCC and cultured in DMEM, 10% FCS, 4 mM L-glutamine, 1 mM sodium pyruvate. MG-132 and FT671 were dissolved in DMSO to prepare a 10 mM stock solution. Cells were treated with either DMSO or the indicated compounds, washed twice with ice-cold PBS and harvested using freshly prepared SDC buffer (room temperature) or urea buffer (chilled to 4°C). The SDC buffer contained 1% SDC, 10 mM TCEP, 40 mM CAA, 75 mM Tris-HCl at pH= 8.5 and its pH was adjusted to 8.5 with 1 N NaOH. The urea buffer contained 8 M urea, 1 mM EDTA, 1 mM CAA, 150 mM NaCl, 50 mM Tris-HCl pH 8.0, 2 µg/ml aprotinin, 10 µg/ml leupeptin, 50 µM PR-619 and 1 mM PMSF, which was added immediately before use. The SDC lysates were heated to 95°C for 10 min while shaking at 750 rpm in a Thermomixer (Eppendorf) and then sonicated for 10 min (10 x 30 sec on/off cycles) using a Bioruptor® Pico sonication device (Diagenode) (for volumes < 2 ml) or a ultrasonic probe device (Bandelin Sonopuls, 1 min, energy output ∼40%, for volumes > 2ml). The urea lysates were processed as described previously^6^. Briefly, the cell pellet was homogenized by pipetting up and down and the lysate cleared by centrifugation at 20,000 x g for 10 min at 4°C. The supernatant was then transferred to a new tube.

### Crosslinking of K-GG antibody

Crosslinking was performed as described previously^6^. In brief, antibody-bound beads were washed with 3 x 1 ml of 100 mM sodium tetraborate (pH 9.0) and then crosslinked by incubating with 1 ml of 20 mM DMP/100mM sodium borate (pH= 9.0) for 30 min at room temperature. The crosslinking buffer was removed and the reaction stopped by washing with 2 x 1 ml of 200 mM ethanolamine (pH= 9.0), followed by incubation with 1 ml of 200 mM ethanolamine (pH= 9.0), with end-over-end rotation at 4°C for 2 h. Finally, the beads were washed with 3 x 1 ml of IP buffer and either directly used or conserved for up to two weeks in 1x phosphate buffered saline (PBS)/0.02% sodium azide.

### K-GG peptide enrichment and LC-MS/MS sample preparation

Protein concentrations were determined using the BCA assay (for experiments in Fig. 1a) or the 660 nm assay (all other experiments, Thermo Fisher Scientific). 2 mg of urea lysate for each replicate was reduced with 5 mM DTT for 45 min and then alkylated with 10 mM CAA for 30 min in the dark before digestion. Proteins were digested with trypsin/Lys-C mix overnight at 37°C (SDC lysates) or at room temperature (urea lysates) with a 1:50 enzyme to protein ratio. For urea samples, the digestion was stopped by adding TFA to a final amount of 0.1% (v/v) and peptides were then desalted using C18 cartridges (Sep-Pak tC18, WAT036790) as follows: a) conditioning with 5 ml of ACN; b) conditioning with 5 ml of 50% ACN/0.1% FA; c) equilibration with 4 x 5 ml of 0.1% TFA; d) loading of the sample, e) washing with 4 x 5 ml 0.1% TFA, f) elution with 2 x 3 ml 50% ACN/0.1% FA. For SDC-lysed samples, the digestion was stopped by adding two volumes of 99% ethylacetate/1% TFA, followed by sonication for 1 min using an ultrasonic probe device (energy output ∼40%). The peptides were desalted using 30 mg (for <1 mg of input, 8B-S029-TAK) or 100 mg (for up to 2 mg of input, 8B-S029-EBJ) Strata-X-C cartridges (Phenomenex) as follows: a) conditioning with 1 ml/3 ml (for 30 mg and 100 mg cartridges, respectively) of isopropanol; b) conditioning with 1 ml/3 ml of 80% ACN/5% NH_4_OH; c) equilibration with 1 ml/3 ml of 99% ethylacetate/1% TFA; d) loading of the sample; e) washing with 2 x 1 ml/3 ml of 99% ethylacetate/1% TFA; f) washing with 1 ml/3 ml of 0.2% TFA; g) elution with 2 x 1 ml/3 ml of 80% ACN/5% NH_4_OH. The eluates were snap-frozen in liquid nitrogen and lyophilized overnight. K-GG peptide enrichment was performed as described with some minor modifications^6^. Briefly, peptides were resuspended in 1 ml of cold immunoprecipitation (IP) buffer (50 mM MOPS pH 7.2, 10 mM Na_2_HPO_4_, 50 mM NaCl) and incubated with 40 µl of a 25 % slurry of cross-linked K-GG antibody-bead conjugate (corresponding to 10 µl beads/IP) for 2 h at 4°C with end-over-end rotation. Beads were then washed twice with 1 ml of IP buffer and an additional time with cold Milli-Q® water. After removing all the supernatant, the beads were incubated with 200 µl of 0.15 % TFA at room temperature while shaking at 1,400 rpm. After briefly spinning, the supernatant was recovered and desalted using in-house prepared, 200 µl two plug StageTips^20^ with SDB-RPS (3M EMPORE™, 2241; for SDC) or C18 (3M EMPORE™, 2215; for urea). SDB-RPS StageTips were conditioned with 60 µl isopropanol, 60 µl 80% ACN/5% NH_4_OH and 100 µl 0.2% TFA. The K-GG enrichment eluate (0.15% TFA) was directly loaded onto the tips followed by two washing steps of 200 µl 0.2% TFA each. Peptides were eluted with 80% ACN/5% NH_4_OH. For C_18_, StageTips were equilibrated with 60 µl ACN and 2 x 60 µl 0.1% FA before sample loading. StageTips were washed with 2 x 60 µl or 0.2% TFA and peptides eluted with 60 µl of 50% ACN/0.1% FA. Peptides were Speedvac dried and then resupended in 10 µl of 0.1% FA, of which 4 µl were injected into the mass spectrometer. For total proteome measurements, a 50 µl aliquot of desalted peptide eluate was transferred to a 0.5 ml tube, Speedvac dried and resuspended in 15 µl of 0.1% FA. The peptide concentration was estimated using a Nanodrop™ device (Thermo Fisher Scientific) and adjusted to 0.4 µg/µl with 0.1% FA, of which 2 µl (800 ng) were injected into the mass spectrometer.

### High pH reversed-phase fractionation

HCT116 cells were treated with MG-132 (10 µM) or FT671 (10 µM) for 6 h before lysis with SDC buffer (see ‘Cell culture, drug treatments and and cell lysis’ for details). 40 mg of each lysate were combined before overnight digestion (37°C) with Trypsin/LysC in a 1:50 enzyme to protein ratio. The resulting peptides were desalted using StrataX-c cartridges (Phenomenex) as described in ‘K-GG peptide enrichment and LC-MS/MS sample preparation’. The lyophilized peptides were resuspended in 0.1 % FA and fractionated using a Zorbax 300SB-C18 column (Agilent Technologies) on an ÄKTA HPLC system (GE Healthcare). Fractionation was performed with a flow rate of 3 ml/min and with a constant flow of 10% 25 mM ammonium bicarbonate, pH 10. Peptides were separated using a linear gradient of ACN from 5% to 35% over 45 min, followed by a 5-min increase to 60% ACN and ramping to 70% over 3 min. Fractions were collected at 60-sec intervals in a 48-well plate to a total of 36 fractions and then pooled to obtain 12 fractions (A1-C1-E1, A2-C2-E2 etc.). All fractions were acidified by addition of FA to a final amount of 0.1% and then lyophilized. Peptides were subsequently resuspended in 1000 µl 0.1% TFA and desalted using Strata-X-C cartridges (Phenomenex) as described. Lyophilized peptides of each fractions were resuspended in 1 ml of IP buffer and enriched for K-GG peptides.

### LC-MS/MS measurements

Peptides were loaded on 40 cm reversed phase columns (75 µm inner diameter, packed in-house with ReproSil-Pur C18-AQ 1.9 µm resin [ReproSil-Pur®, Dr. Maisch GmbH]). The column temperature was maintained at 60°C using a column oven. An EASY-nLC 1200 system (ThermoFisher) was directly coupled online with the mass spectrometer (Q Exactive HF-X, ThermoFisher) via a nano-electrospray source, and peptides were separated with a binary buffer system of buffer A (0.1% formic acid (FA) plus 5% DMSO) and buffer B (80% acetonitrile plus 0.1% FA plus 5% DMSO), at a flow rate of 250 nl/min (for 75 min and 125 min gradients). For the 60 min method, a flow rate of 300 nl/min was used. For the 30 min and 45 min gradients, a flow rate of 350 nl/min was used and for the 15 min DIA method, a 15 cm column and a flow rate of 500 nl/min were used. The mass spectrometer was operated in positive polarity mode with a capillary temperature of 275°C.

The DDA method consisted of a MS1 scan (m/z= 300-1,650, R= 60,000, maximum injection time= 20 ms, spectrum data type= profile) followed by TopN MS/MS scans (N= 15). These were acquired at R= 15,000, AGC target = 1e5, maximum injection time= 28 ms, isolation window 1.4 Th, NCE= 27 and a scan range of 200-2000 m/z.

The DIA methods consisted of a MS1 scan (m/z= 300-1,650) with an AGC target of 3×10^^6^ and a maximum injection time of 60 ms (R= 120,000). DIA scans were acquired at R= 30,000, with an AGC target of 3×10^^6^, ‘auto’ for injection time and a default charge state of 4. The spectra were recorded in profile mode and the stepped collision energy was 10% at 25%. The number of DIA segments was adjusted to achieve an average of 4 data points per peak.

### Raw data processing

MS raw files acquired with data-dependent acquisition mode were analyzed using MaxQuant^11^ version 1.6.10.43 (maxquant.org), data-independent acquisitions with DIA-NN^10^ version 1.7.10 (https://github.com/vdemichev/DiaNN). Reviewed UniProt entries (SwissProt, release 10-2018, 42,427 entries) were used as protein sequence database for both MaxQuant and DIA-NN searches. For MaxQuant searches, default settings were used (enzyme = trypsin/P, two missed cleavages, carbamylation of cysteines as fixed modification and protein N-terminus acetylation and oxidation of methionine as variable modifications) and K-GG was added as variable modification. ‘Match between runs’ was enabled were indicated. For DIA-NN, one missed cleavage, a maximum of two variable modifications and N-terminal excision of methionine were allowed. Carbamylation of cysteines was set as fixed modification, oxidation of methionine and K-GG (UniMod: 121) as variable modification in case of ubiquitinomics. The quantification strategy was set to ‘robust LC (high precision)’. For library-free searches, the mass accuracy was set to 30 ppm and ‘deep learning-based spectra and RTs prediction’ was enabled only in case the proteome. For ubiquitinomics data, a training library was included in the search instead (where indicated). This training library was created in DIA-NN by searching the respective DIA files with the library-free option and ‘deep learning-based spectra and RTs prediction’ disabled. All library-free searches were done in two steps: first, a spectral library was created in a library-free search using all DIA raw files of interest (‘deep learning-based spectra and RTs prediction’ disabled). Then, the resulting spectral library was used in a regular library-based search. For data shown in Fig. 2, the DIA-NN q-value filter was set to 1.0 to obtain an almost complete data matrix (i.e. only precursors quantities are missing, for which no peak could be picked).

### Bioinformatic data analysis

In MaxQuant outputs (DDA), ubiquitinated peptides (K-GG) were filtered for reverse hits, contaminants and non-K-GG peptides and counted using the Perseus software^21^. For DIA-NN outputs in Figure 1, precursors were aggregated to peptides by taking the mean intensities and K-GG peptides (UniMod: 121) were counted. Coefficients of variation were computed on the K-GG intensities and the plots were created with GraphPad Prism 8 (GraphPad Software). The GO term enrichment analysis was done with Perseus^21^. DIA-NN outputs of Fig. 2 were further processed with R (version 3.6.1). The DIA-NN precursor q-value filter was set 1.0, to obtain almost complete data matrices for both proteome and ubiquitinome data sets (missing only data of precursors where no peak could be picked).

The ubiquitinomics precursor data matrix were then filtered based on q-values according to the following rules: precursors were not excluded, if they pass the q-value< 0.01 threshold in a) at least 50% of all samples, or b) in 100% of samples of at least one experimental condition. K-GG precursor intensities were aggregated to peptides by the MaxLFQ^22^ algorithm, as implemented in the DIA-NN R package (https://github.com/vdemichev/diann-rpackage/). Instead of using the protein groups created by DIA-NN, protein inference was performed in-house following the ID Picker^23^ algorithm. K-GG peptide to site mapping was done using reviewed entries of the human UniProt database (SwissProt, release 10-2018, 42,427 entries). The peptide intensities were normalized by median scaling before differential expression analysis.

The precursor matrices of the proteomes were filtered for precursors that pass the <0.01 q-value filter in a) 50% of samples or b) at least 80% of all DMSO control samples. Protein intensity normalization was done using the MaxLFQ algorithm^22^.

Significance testing was performed with LIMMA^24^, where FT671 treatments were compared to their corresponding DMSO controls at each time point. For both ubiquitinome and proteome data sets, a batch factor was added to the LIMMA model to account for the two batches during sample processing (Figure 2). Multiple testing corrected q-values <0.05 were considered as significant. For comparing proteome and ubiquitinome data, protein groups of each dataset were disaggregated into individual UniProt identifiers and then mapped between the data sets. For individual plots, the protein groups of the ubiquitinome data sets along with K-GG site positions were used. The effective FDR of the high pH reversed-phase fractionated K-GG peptide library (built by library-free search in DIA-NN) was verified by using a two-species empirical FDR approach, as established by Bruderer et al^9^ and adapted by Demichev et al^10^.

Cytoscape (version 3.7.2, cytoscape.org) was used for mapping the combined interactomics data from BioGrid^19^ with the ubiquitinomics data from this study. BioGrid interaction data (version 3.5.186) was filtered for human proteins with physical interactions with USP7 in at least four studies. Ubiquitination sites were mapped onto Usp7 and its interactors, if they showed >2-fold significant upregulation at 15 minutes. Proteins without significant ubiquitination regulation are not shown.

## Supplementary Materials

Supplementary Figures 1-9

Supplementary Table 1-6

## Figure Legend

**Supplementary Fig. 1.**
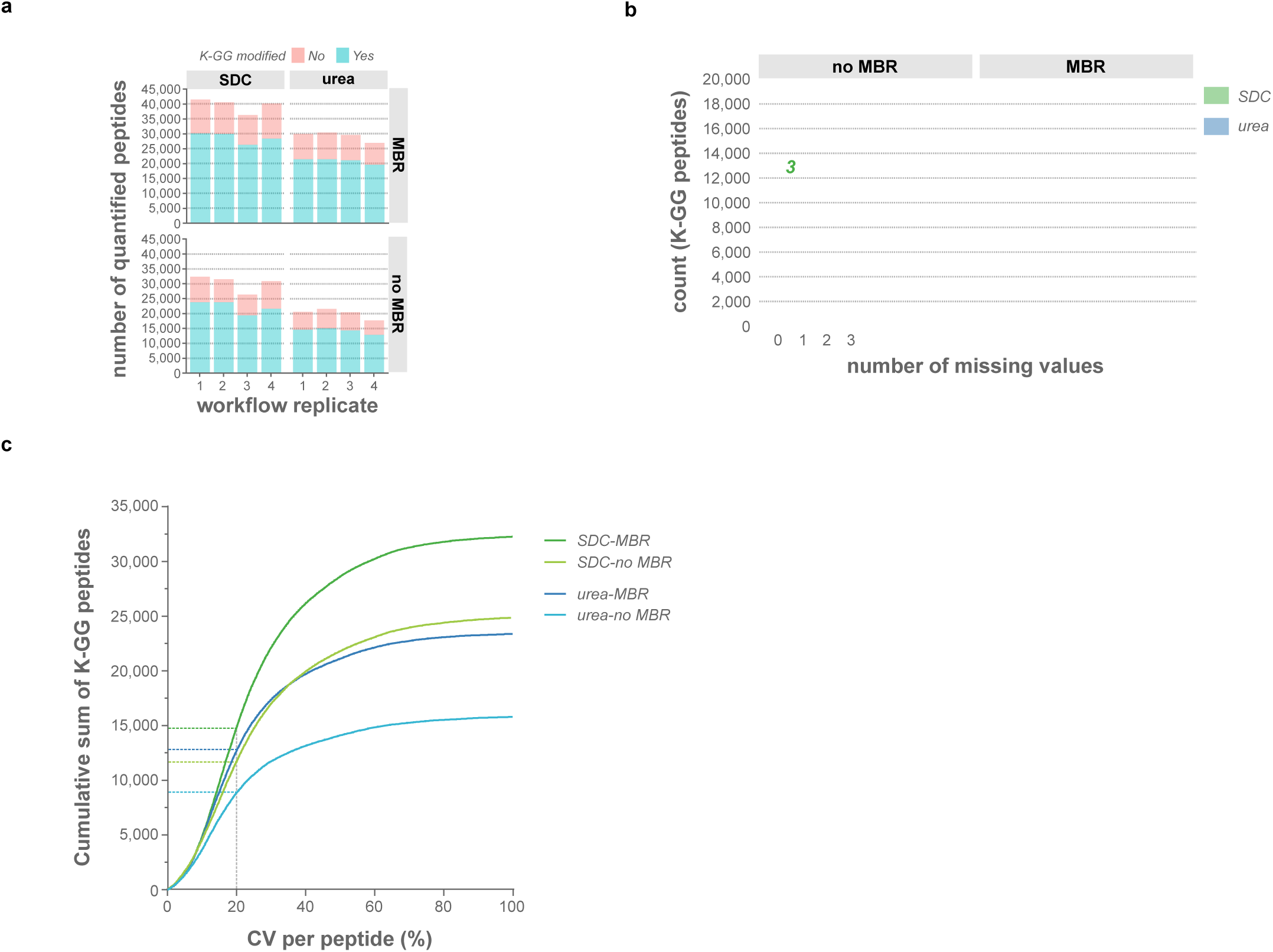
Side-by-side comparison of urea- and SDC-based lysis protocols for ubiquitinomics. **a**, Fraction of unmodified and K-GG modified peptides quantified from either urea or SDC lysates in HCT116 cells. Four individual samples were processed for each lysis protocol, with 2 mg of protein input per replicate. MS samples were analyzed in DDA mode with a 125 min gradient MS method, followed by data processing with MaxQuant. **b**, Number of K-GG peptide identifications with 0, 1, 2, or 3 missing values for the SDC (green) and urea (blue) lysis methods. Numbers on top of the bars show cumulative K-GG peptide IDs across all four replicates analyzed for each lysis protocol. **c**, Ranked K-GG peptide coefficients of variation (CVs) for SDC- and urea-based lysis, with or without the match between run (MBR) option enabled in MaxQuant. The 20% CV cut-off is marked.

**Supplementary Fig. 2.**
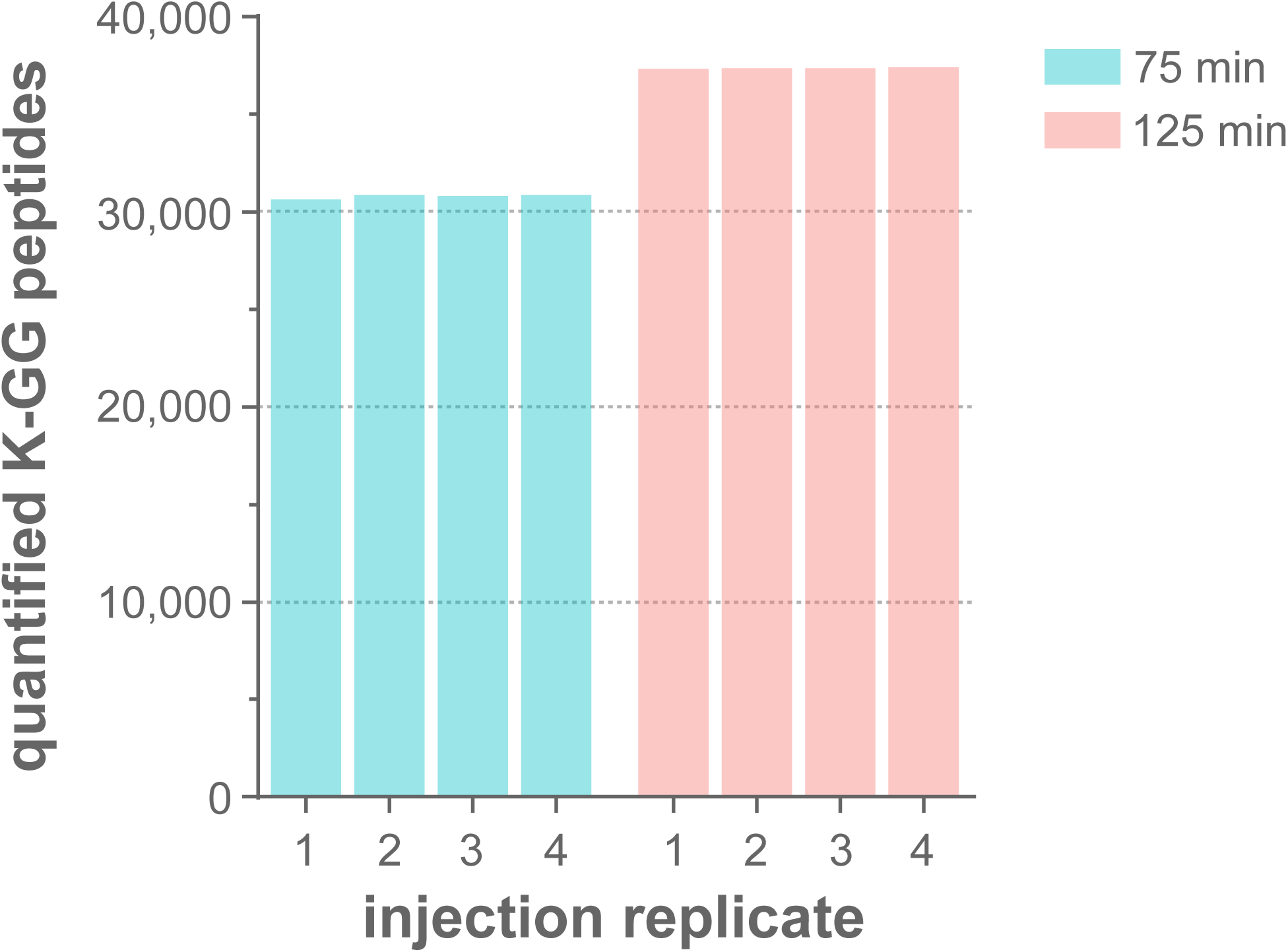
Comparison of 75 min and 125 min DIA-MS methods. Upon K-GG remnant peptide enrichment from HCT116 SDC lysates, K-GG peptides were analyzed by DIA with 75 min and 125 min LC-MS methods. The data was analyzed with DIA-NN featured library-free search option without deep learning (see *Methods*).

**Supplementary Fig. 3.**
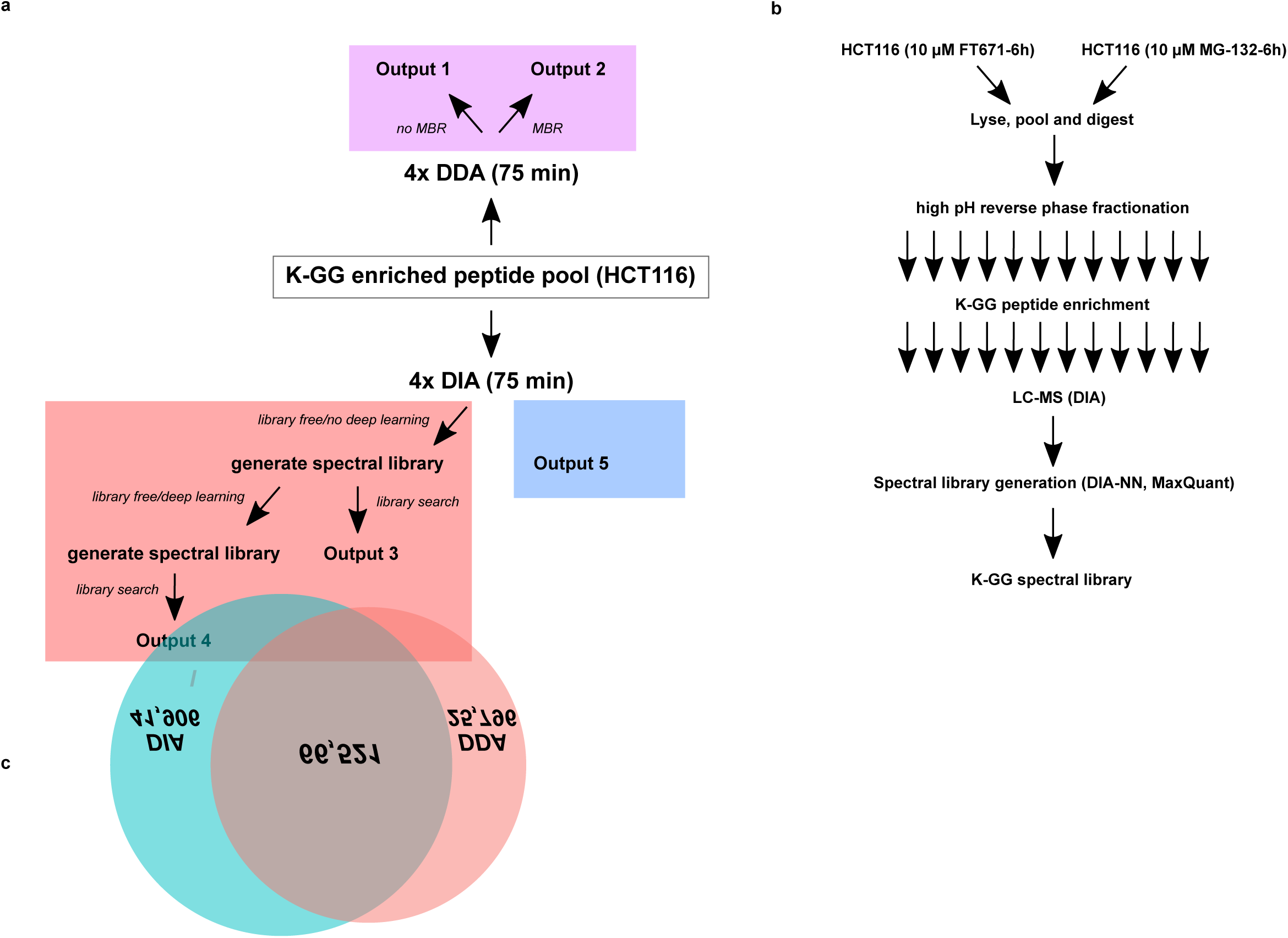
Direct comparison of DDA and DIA for ubiquitinomics. **a**, Schematic of data-dependent acquisition (DDA) and data-independent acquisition (DIA) strategies showing the different search options applied for both MS acquisition modes. Pooled K-GG peptide sample material was prepared from MG-132-treated HCT116 cells and four replicates were analyzed by either DDA or DIA MS. The DDA data was processed with MaxQuant, with or without ‘match between runs (MBR)’ enabled, creating two outputs. For DIA data processing in DIA-NN, the same DIA MS raw files were used to explore both library-free (pink rectangle) and library-based (blue rectangle) search options. For library-free searches, a spectral library was first created with ‘deep learning-based spectra and RTs prediction’ disabled. This library was then used in a library-based search to create output 3. The same library was included as training library in a second library-free search (‘deep learning-based spectra and RTs prediction’ disabled), thus creating a second spectral library. The resulting library was then used in another library-based search to create output 4. A spectral library of K-GG enriched peptides was created by high pH reversed-phase fractionation and used in a regular library-based search to create output 5. **b**, For ultra-deep K-GG peptide spectral library generation, HCT116 cells were treated with either MG-132 or FT671 for 6 h (10 µM each) before lysis in SDC buffer. 40 mg of each lysate were combined and digested with trypsin. The peptides were fractionated into 12 fractions using high pH reversed-phase fractionation and the individual fractions were enriched for K-GG peptides. Each fraction was measured with both DIA (library-free search, with ‘deep learning-based spectra and RTs prediction’ disabled) and DDA mode. DIA-NN and MaxQuant were used for data analysis, respectively. **c**, Venn diagram of ubiquitinated peptides identified by DIA and DDA using the fractionation and K-GG enrichment workflow shown in **b**.

**Supplementary Fig. 4.**
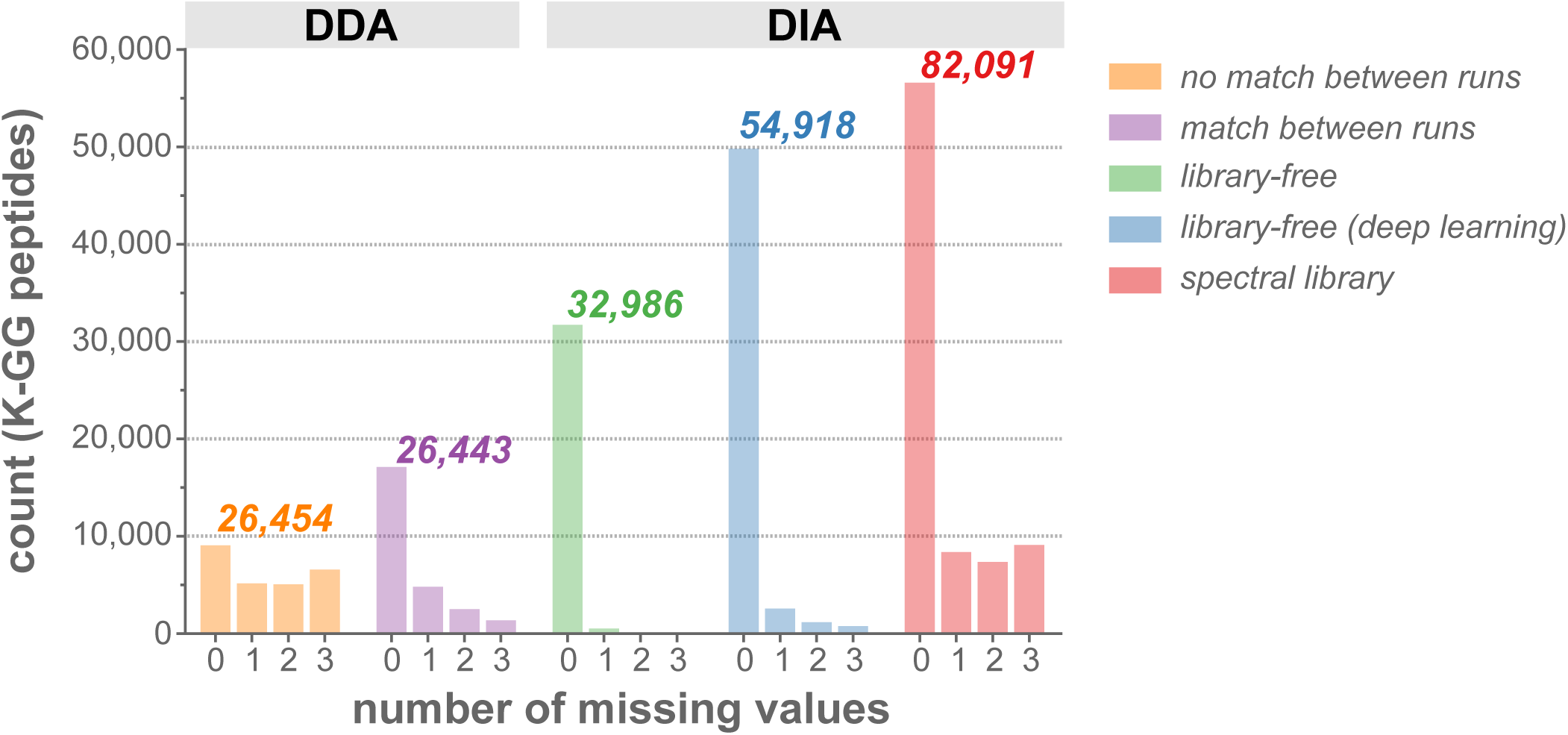
Comparison of DIA and DDA for single-shot ubiquitinomics. Numbers of identifications with 0, 1, 2, or 3 missing values in four replicates are shown for different search options of DDA and DIA. Numbers on top of the bars represent the cumulative K-GG peptide IDs for the different data acquisition and processing methods tested.

**Supplementary Fig. 5.**
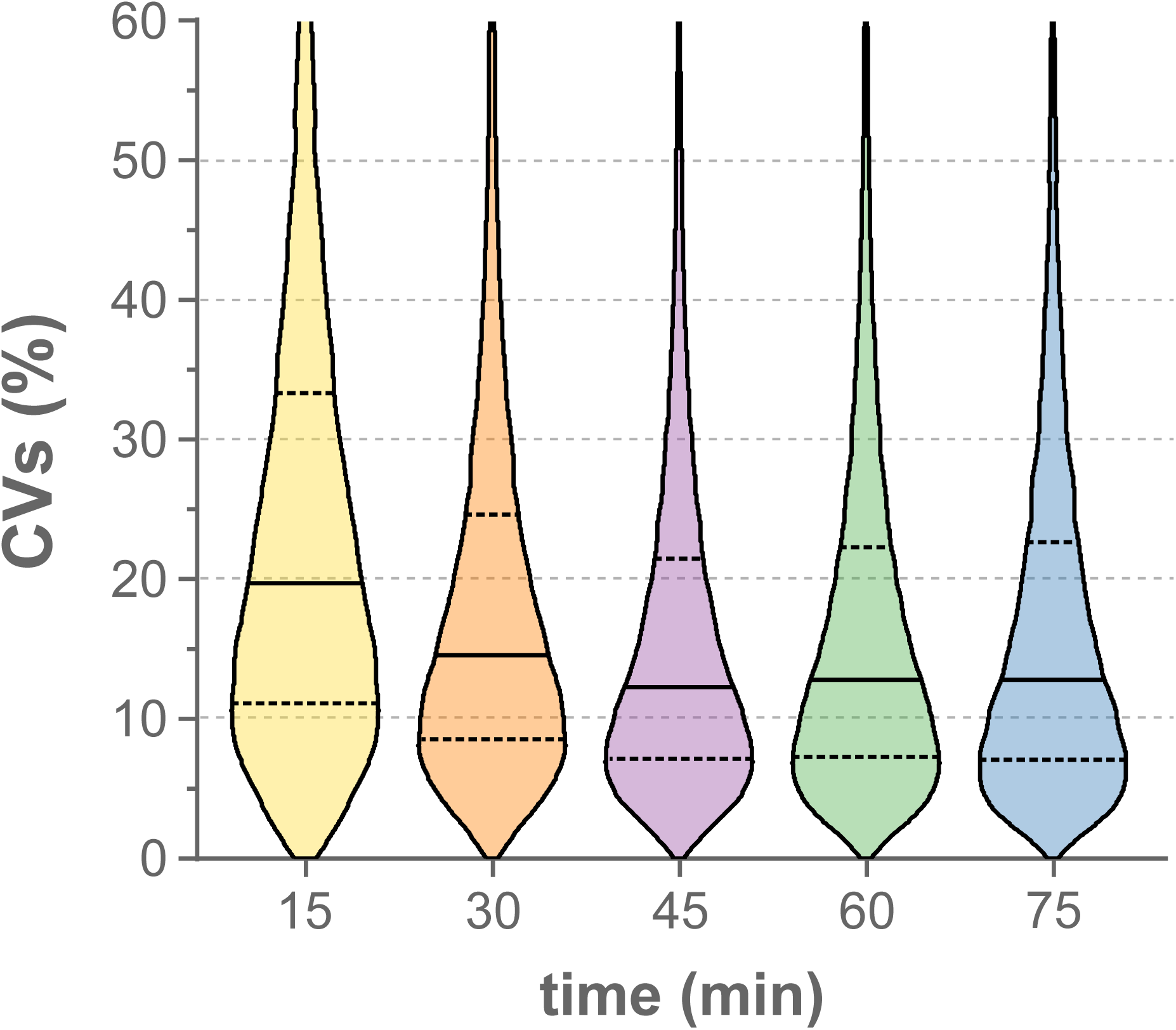
Evaluation of shorter LC gradient lengths to increase throughput in single-shot DIA-MS ubiquitinomics. Coefficients of variation (CVs) distributions for K-GG peptides quantified in four replicates in 15, 30, 45, 60 and 75 min DIA-MS single-shot runs. DIA data were processed with the ultra-deep K-GG peptide spectral library (see *Methods* and Supplementary Fig. 5b for details). Continuous lines in the violin plots demarcate the median and dashed lines upper and lower quartiles.

**Supplementary Fig. 6.**
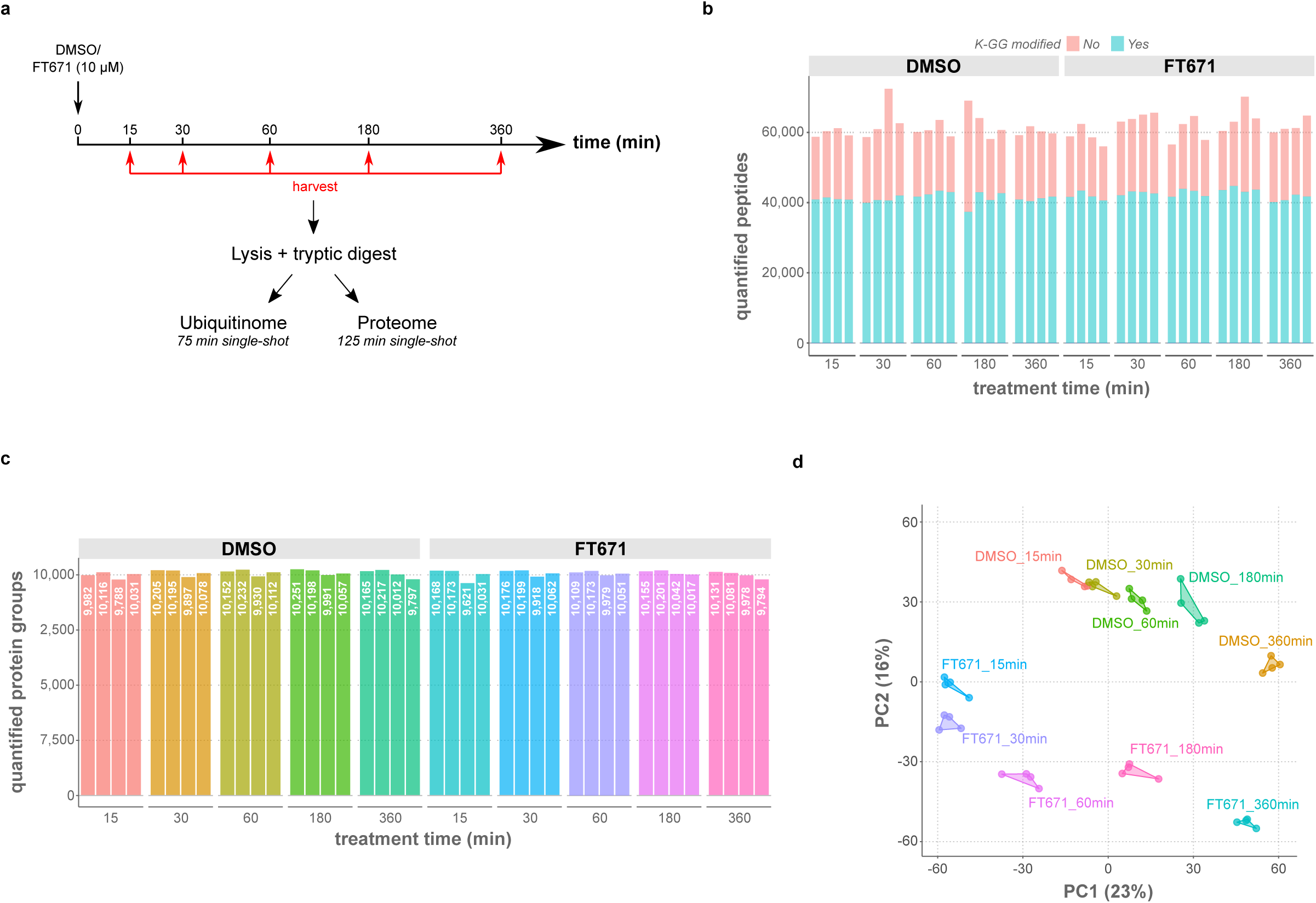
FT671 time course experiment to measure USP7-mediated changes in the ubiquitinome and the proteome. **a**, Schematic of the FT671 time course experiment. HCT116 cells were treated with 10 µM of FT671 or DMSO and harvested in SDC buffer at the indicated time points. After tryptic digestion of 2 mg of total protein per sample, both the ubiquitinome and the proteome were quantified in single-shot DIA-MS runs using DIA-NN for spectral library-based searches. **b** and **c**, Numbers of MS-quantified K-GG peptides and proteins at 1% FDR, shown for all replicates and treatment conditions. **d**, Principal component analysis of the ubiquitinomics data.

**Supplementary Fig. 7.**
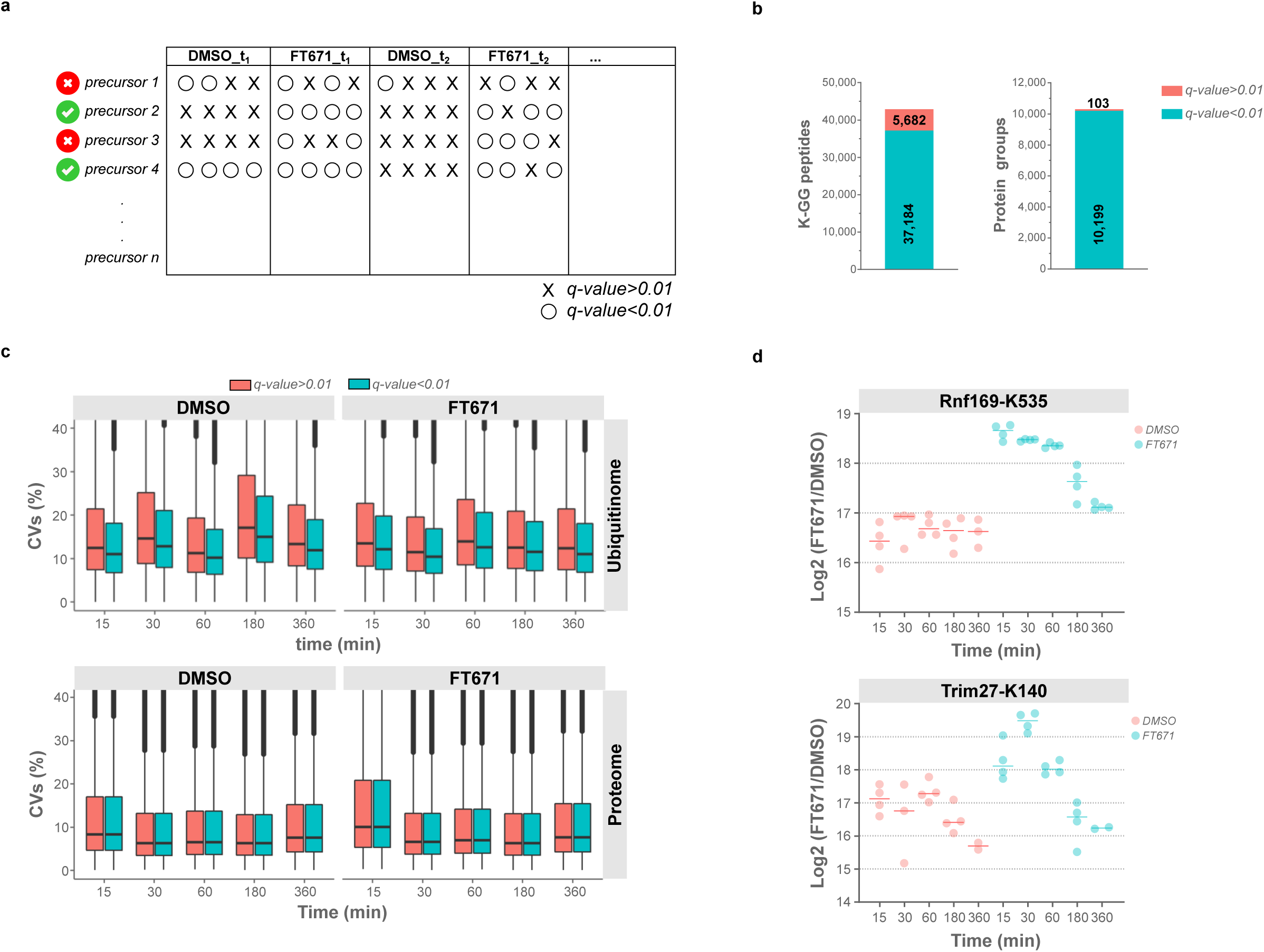
Strategy to minimize the number of missing values in DIA-MS data. **a**, To obtain ubiquitinomics and proteomics data matrices without missing values, the precursor q-values were set to 1. The resulting data matrices were then filtered for identifications that had q-values < 0.01 in 50% of all samples, or in all four replicate samples of at least one experimental condition (ubiquitinome). These rules were selected to remove incomplete quantifications across all samples and at the same time keep K-GG peptide identifications of potential primary enzyme targets, which were below the detection limit in DMSO controls and induced upon USP7 inhibition. In case of the proteome, the filtering rules were as follows: q-value< 0.01 in 50% of all samples or in 80% across all DMSO controls. The latter rule was chosen to retain precursor quantifications of USP7-ubiquitinated proteins that are degraded at later time points. **b**, Number of K-GG peptides and proteins that are additionally quantified when including low-confidence precursor identifications (q-value> 0.01) to increase data completeness, as described in (**a**). **c**, The complete data matrices of the ubiquitinome and the proteome were filtered based on the q-value as described in (**a**). Shown are coefficients of variation (CVs) of all quantified K-GG peptides or proteins obtained according to the filtering rules detailed in (**a**), either including or excluding identifications, for which the q-value was > 0.01 in at least one sample. **d**, Selected K-GG sites of known Usp7 substrates, which would have been excluded from statistical testing because of missing values. By replacing missing values with quantitative data from low-confidence identifications, both sites were identified as significantly upregulated upon Usp7 inhibition.

**Supplementary Fig. 8.**
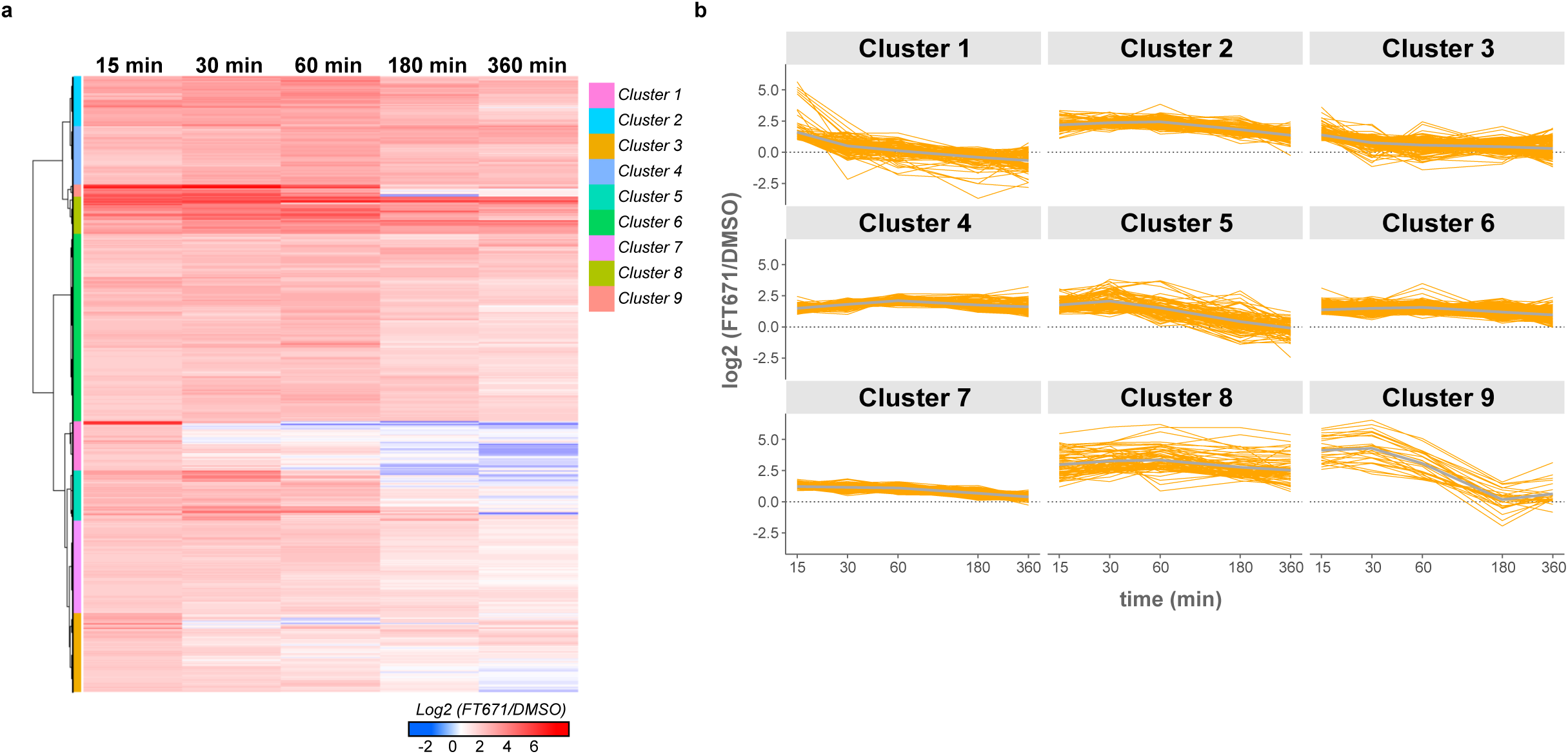
Temporal profiles of FT671-regulated K-GG peptides. **a**, Hierarchical clustering of the temporal profiles of all ubiquitinated peptides that were significantly upregulated by more than two-fold upon 15 min FT671 treatment. Nine distinct clusters were identified for this subset of peptides that underwent rapid ubiquitination after USP7 inhibition. **b**, Profiles of K-GG peptides assigned to the different clusters are shown. Average profiles are highlighted in grey. Most K-GG sites are maximally induced at 15 min and either decrease in intensity at later time points or remain unchanged throughout the time course.

**Supplementary Fig. 9.**
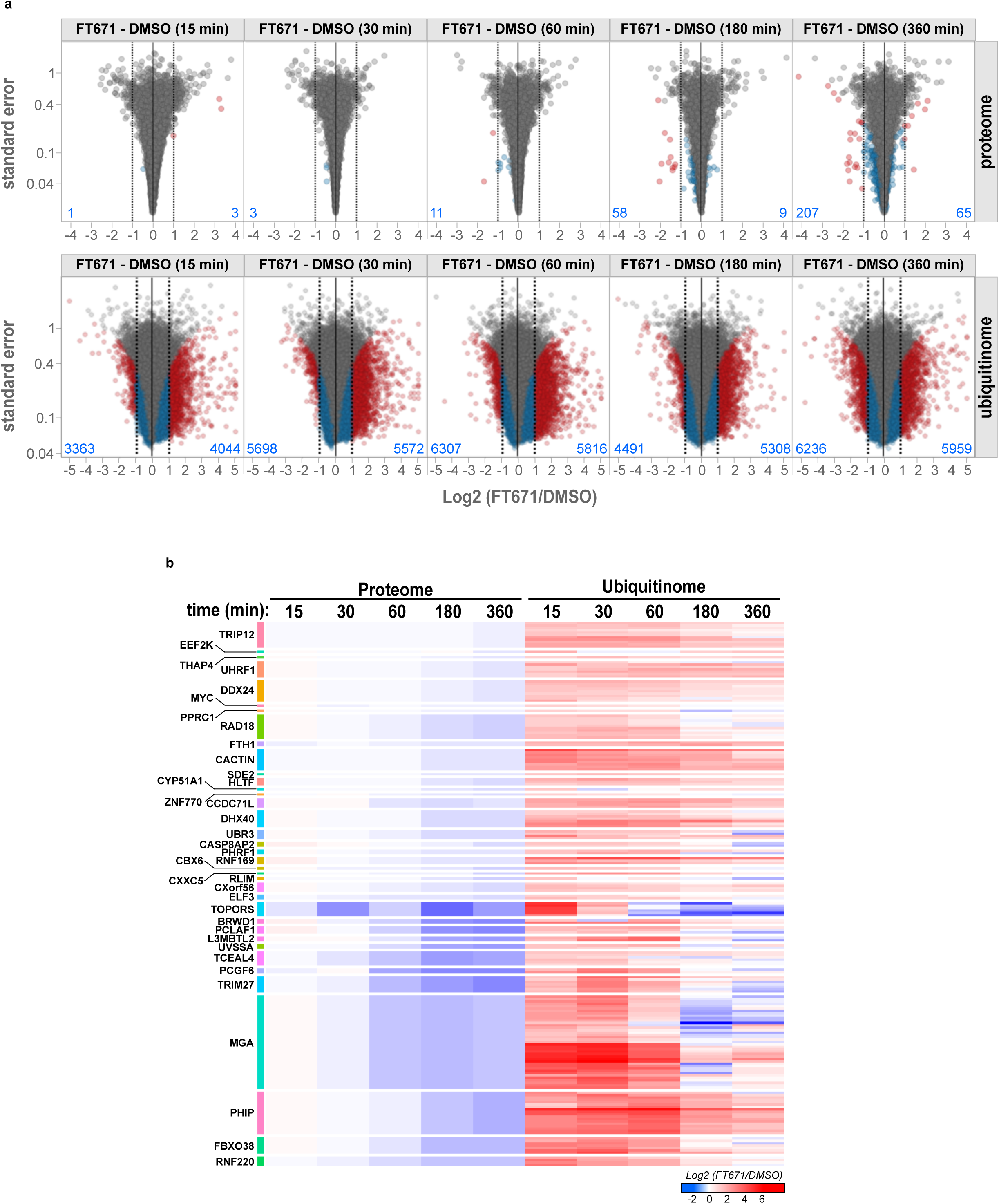
USP7-dependent changes in the ubiquitinome and the proteome. **a**, Volcano plot visualizations of log_2_-transformed, average FT671/DMSO ratios versus their standard errors, as determined by quantitative DIA-MS analysis of proteins (upper panels) and ubiquitinated peptides (lower panels). Significantly regulated proteins and K-GG peptides are colored (fold changes <2 in blue and fold changes >2 in red) and the number of significant up- and downregulations at 5% FDR are indicated for each volcano plot. **b**, Heat map of proteins that were significantly downregulated by more than 20% in the FT671 time course experiment with significant, and more than two-fold induction of at least one ubiquitination site at 15 min.

